# Recovering false negatives in CRISPR fitness screens with JLOE

**DOI:** 10.1101/2022.03.22.485359

**Authors:** Merve Dede, Traver Hart

**Affiliations:** Department of Bioinformatics and Computational Biology, The University of Texas MD Anderson Cancer Center, Houston, TX; Department of Cancer Biology, The University of Texas MD Anderson Cancer Center, Houston, TX

## Abstract

It is widely accepted that pooled library CRISPR knockout screens offer greater sensitivity and specificity than prior technologies in detecting genes whose disruption leads to fitness defects, a critical step in identifying candidate cancer targets. However, the assumption that CRISPR screens are saturating has been largely untested. Through integrated analysis of screen data in cancer cell lines generated by the Cancer Dependency Map, we show that a typical CRISPR screen has a ∼20% false negative rate, in addition to library-specific false negatives. Replicability falls sharply as gene expression decreases, while cancer subtype-specific genes within a tissue show distinct profiles compared to false negatives. Cumulative analyses across tissues improves our understanding of core essential genes and suggest only a small number of lineage-specific essential genes, enriched for transcription factors that define pathways of tissue differentiation. To recover false negatives, we introduce a method, Joint Log Odds of Essentiality (JLOE), which builds on our prior work with BAGEL to selectively rescue the false negatives without an increased false discovery rate.

## Introduction

The search for essential genes - genes whose loss of function results in a severe fitness defect - has been of outstanding interest to the scientific community. Prior to advanced genomic technologies, the assumption was that the majority of genes were essential for life (Horowitz & Leupold, 1951). This idea was disproven by several studies that utilized saturating random mutagenesis to show that in *C. elegans* and *S. cerevisiae*, 12-15% of the genome was estimated to be essential (Brenner, 1974; Goebl & Petes, 1986). These studies were limited by the methods at the time and the lack of the availability of complete genome sequences.

With the advances in genome technologies that enabled sequencing of eukaryotic organisms, systematic gene knockout studies were performed in *S. cerevisiae*, identifying essential genes by deletion of open reading frames in the yeast genome (Giaever et al., 2002; Winzeler et al., 1999). This method identified 17% of yeast genes as essential for growth in rich medium (Winzeler et al., 1999). Generation of siRNA and shRNA libraries to conduct genome-wide RNAi screens facilitated the study of essential genes in other model organisms (Dietzl et al., 2007; Kamath et al., 2003; Meister & Tuschl, 2004; Moffat & Sabatini, 2006). In these RNAi screens, 30% of the genome was shown to be essential in *D.melanogaster* cell lines, (Dietzl et al., 2007), compared to only 8.5% of the *C.elegans* genome in whole worms (Kamath et al., 2003). A later study showed that a binary classification of genes into essential and non-essential was misleading due to the context dependent nature of gene essentiality and that 97% of yeast genes showed some growth phenotype under different environmental conditions (Hillenmeyer et al., 2008).

Identifying essential genes in human cancer cells is of special interest in oncology since the cancer-specific essential genes represent genomic vulnerabilities that can potentially be targeted with novel therapeutic agents. Studies using CRISPR/Cas9 technology revealed that mammalian cells have more essential genes than RNAi screens were able to detect and that, at the same false discovery rate, CRISPR screens generated 3-4 times more essential genes (T. Hart et al., 2014). Moreover, multiple groups revealed lists of ∼2000 highly concordant human essential genes, and comparison of CRISPR technology to orthogonal techniques such as random insertion of gene traps also showed consistent results (Blomen et al., 2015; T. Hart et al., 2015; T. Wang et al., 2015). These findings were initially thought to indicate that the CRISPR-Cas9 screens are near-saturating and that a well-designed screen can detect a cell’s full complement of essential genes. However, it is still poorly understood whether possible systematic biases in CRISPR screens affect our understanding of human cell-intrinsic gene essentiality. In the absence of a ground truth, the actual true positive, false positive and false negative rates of a typical genome-wide CRISPR-Cas9 knockout screen are still unknown.

In this study, we examine some of the biases characteristic of genome-wide CRISPR-Cas9 knockout screening. Using publicly available genome wide screen data from 769 genetically heterogeneous cell lines from the Cancer Dependency Map initiative (Meyers et al., 2017; Tsherniak et al., 2017), we demonstrate the systematic biases in these screens, investigate and model the actual number of essential genes identifiable with CRISPR technology, estimate the false discovery rate (FDR) and false negative rate (FNR) in a CRISPR screen, and describe a computational method, the Joint Log Odds of Essentiality (JLOE), that can rescue these false negatives. We show that JLOE improves screen concordance with the Sanger dataset (Behan et al., 2019) and reduces the number of background-specific essential genes.

## Results

To systematically evaluate the biases in genome-wide CRISPR-Cas9 knockout screening, we processed the raw read counts of loss of function screens performed in 769 genetically heterogeneous cell lines from the 2020Q2 release of the publicly available Avana data (Meyers et al., 2017). We applied our computational pipeline as described in the methods section to correct for copy number effects using the previously described CRISPRcleanR (Iorio et al., 2018) and BAGEL (T. Hart & Moffat, 2016; E. Kim & Hart, 2021) algorithms to assign an essentiality score (Bayes Factor, BF) to each gene in every screen. After applying quality control metrics (see Methods), our final dataset included 659 screens meeting the F-measure criteria of 0.80 and above (**Supplementary Figure 1A-B, Supplementary Table1**). These 659 cell lines are derived from 25 tissue types, with varying representation (**Figure 1A**).

**Figure 1.**
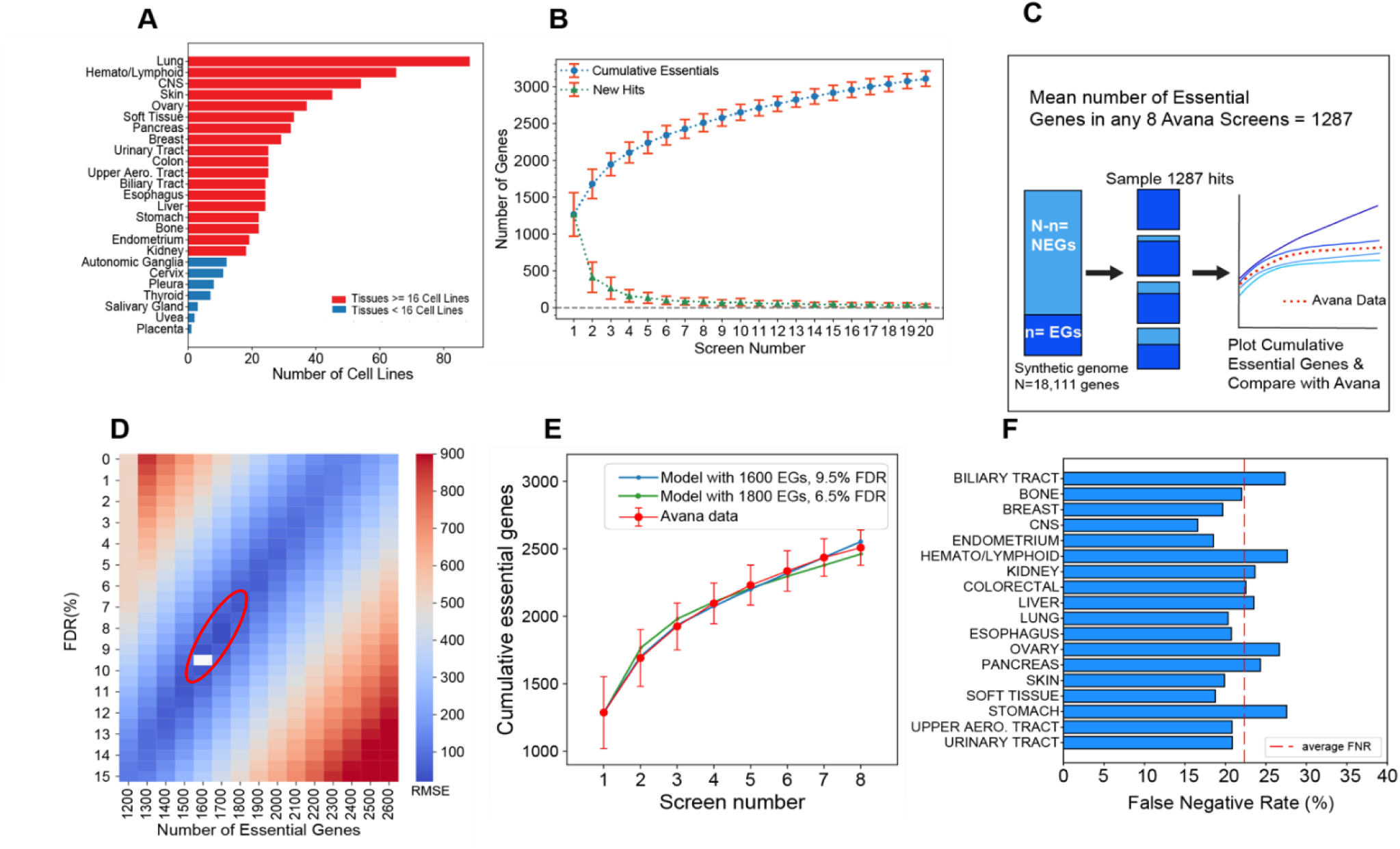
Synthetic genome modeling. **A)** The representation of each tissue type in the 659 high-performing screens in the Avana data. Tissue types with >=16 high-performing cell lines are indicated with red bars. **B)** The cumulative number of essential genes (BF>=10) and newly discovered essential gene curves in any randomly selected set of 20 Avana screens show a curve that converges into a positive slope. Blue circles indicate the number of cumulative essential genes in each screen and the green triangles show the number of newly discovered essential gene hits in each consecutive screen. The error bars represent the standard deviation of the number of cumulative essential genes and the newly discovered essentials in 100 iterations. **C)** Diagram overview of the synthetic genome modeling of essential genes. From a synthetic genome of 18,111 genes, 1287 hits were randomly sampled from the essential (n) and non-essential gene (NEG) populations based on the defined false discovery rate (FDR) in the simulation. The resulting cumulative essential hits across 8 iterations were plotted and compared to the mean cumulative essentials curve from the Avana data. **D)**The synthetic genome modeling reveals the best fitting model with n=1600 genes and 9.5% FDR (white box). **E)** Plot showing the model with the best fit (in blue) to the Avana data (in red). **F)** Average screen false negative rate (FNR) as determined by comparing the number of essential genes from the best fit model in each tissue type to the mean number of observed essentials in each tissue type. The red dashed line indicates a mean FNR of 21.2% across all tissue types tested.

### Synthetic genome modeling of essential genes in 659 cell lines in the Avana data

To estimate the total number of essential genes in a cell, we employed a method based on the cumulative essentiality observations across all screens in the Avana data. Our approach is based on the expectation that, for a sufficient number of identical screens without any false positives, a plot of the cumulative number of essential genes would flatten to a slope of zero when all of the essential genes in the population are identified. In contrast, in a population of screens with either cellular heterogeneity or some degree of false discovery rate (or most likely both), the slope of the cumulative essential genes plot would remain positive, reflecting the continuous accumulation of false positives as well as cell line specific essential genes in otherwise saturated screens. This principle was previously demonstrated to estimate the total population of essential genes detected in shRNA screens (T. Hart et al., 2014).

We applied this logic to the Avana dataset and plotted the cumulative essential genes across sets of 20 cell lines that are randomly selected without replacement from all Avana screens (**Figure 1B**). The sampling process was repeated 100 times to minimize bias in the selection of screens. To define gene essentiality, a strict threshold of BF>10 was used for every gene in each screen since this threshold represents a posterior probability of gene essentiality of ∼99% **(Supplementary figure 1C)**. Using this BF 10 threshold yielded a cumulative essentials curve that converged to a positive slope, which was consistent with previous observations in the shRNA screens **(Figure 1B**).

We propose a model of cell-intrinsic gene essentiality and CRISPR screen sensitivity that describes the trend shown in Figure 1B. First, there exists a large population of essential genes across all screens. Second, we posit the existence of some saturation point, here shown at k=8 screens (n = ∼2,500 cumulative essential genes). Given this saturation point, it follows that all essential genes are never captured in a single experiment and each screen carries some unknown false negative rate (FNR). Multiple screens are needed before all essential genes in the population can be identified. Third, after the saturation point is reached, additional screens continue to accumulate a combination of false positives as well as context-specific essential genes that were either not detectable or not present in the prior set of screens, and that the rate at which these genes are observed offers some estimate of the false discovery rate (FDR) of each screen.

To estimate the parameters of this model, we carried out repeated screens *in silico* and compared synthetic cumulative essential curves to those derived from the data. Starting with a genome of N=18,111 genes – the number of genes tested in the Avana library – we arbitrarily defined *n* essential genes, leaving N-*n* nonessential. We further defined a screen false discovery rate hyperparameter between 1-15%. Then we repeatedly sampled this genome with a screen that randomly drew 1,287 hits – the mean number of hits at BF>10 in a single Avana screen; **Figure 1B** – from the essential and nonessential populations based on the FDR hyperparameter (e.g. at 10% FDR, 129 nonessentials and 1,158 essentials were randomly selected; see Error! Reference source not found.**C)**. Finally, we determined the cumulative hits across k=8 replicate screens, estimating that eight samples was a good estimate of screen saturation in the data (**Figure 1B**) and judging that it was more important to fit the model to our observations in this region than in the saturated region (k>>8). We calculated the root-mean-squared error from the mean cumulative essentials curve determined from the Avana data and plotted RMSE vs. the two model hyperparameters **(Figure 1D)** observing that the best fit occurred with *n*=1,600 essential genes and FDR=9.5% **(Figure 1E)**. Notably, a region of good fits, with RMSE < 2xRMSE_min_, occurs between *n=*1,500-1,800 essential genes and a corresponding decrease in per-screen FDR from 9.5% to ∼6.5% **(Supplementary Figure 1).**

Given the broad range of lineages from whence the screened cell line models were derived, it is clear that some of the context-specific essential genes will be incorrectly classified as false positives in our synthetic genome modeling approach. In order to minimize the effect of these tissue-specific essential genes, we repeated the analysis using cell lines from only one tissue type, filtering for 18 tissues that are represented by at least 16 high quality screens **(Figure 1A)**. Each tissue type yielded similar cumulative essential curve as shown here for the colorectal cancer cell lines in the Avana data (**Supplementary Figure 2)**. We repeated this modeling approach in each tissue type and obtained remarkably similar results (**Supplementary Figures 3 and 4)**. Next, we compared the best-fit number of essential genes from the synthetic genome model in each tissue type to the mean number of essential gene hits observed in that same tissue type and calculated an average false negative rate across each tissue **(Figure 1F**). Across all tissue types we examined, we determined the mean FNR to be ∼20% in each screen.

### Saturation modeling to differentiate essential genes and false positives

While our synthetic genome approach gave us an estimate of the expected total number of essential genes and the FDR in an average screen, it does not provide any way to differentiate true hits from false positives. To address this issue, we utilized an alternative view of the saturating behavior of CRISPR screens, similar to the methodology described for shRNA screens (T. Hart et al., 2014). Based on our judgment that screening in virtually all lineages achieved saturation after approximately eight cell lines, we again selected the lineages with at least twice this number of screens indicated in **Figure 1A**. From each lineage, we randomly selected eight screens (“initial screens”) without replacement, identified the cumulative number of essential genes (defined by BF>10) and determined the number of cell lines in which each gene was classified as essential. We then randomly selected an additional eight screens (“subsequent screens”), again without replacement, and determined the number of cell lines in which each gene was classified as new hits – that is, BF>10 but not a hit in any of the initial eight screens (**Figure 2A)**. We assumed that all of these new hits were false positives, and that the histogram of observations of these false positives estimates the frequency of false positives in the initial screens. It is almost certainly not the case that all of these are actually false positives, given the known presence of tumor subtypes within each tissue/lineage, the high likelihood of subtype-specific essential genes, and the probability that any given subtype escaped being selected in the initial eight screens. However, this assumption is useful for modeling purposes, as it provides an estimate of the upper bound of the false discovery rate using this saturation modeling approach. We repeated this process 100 times and plot the resulting distributions of hits vs. number of screens **(Figure 2A)**.

**Figure 2.**
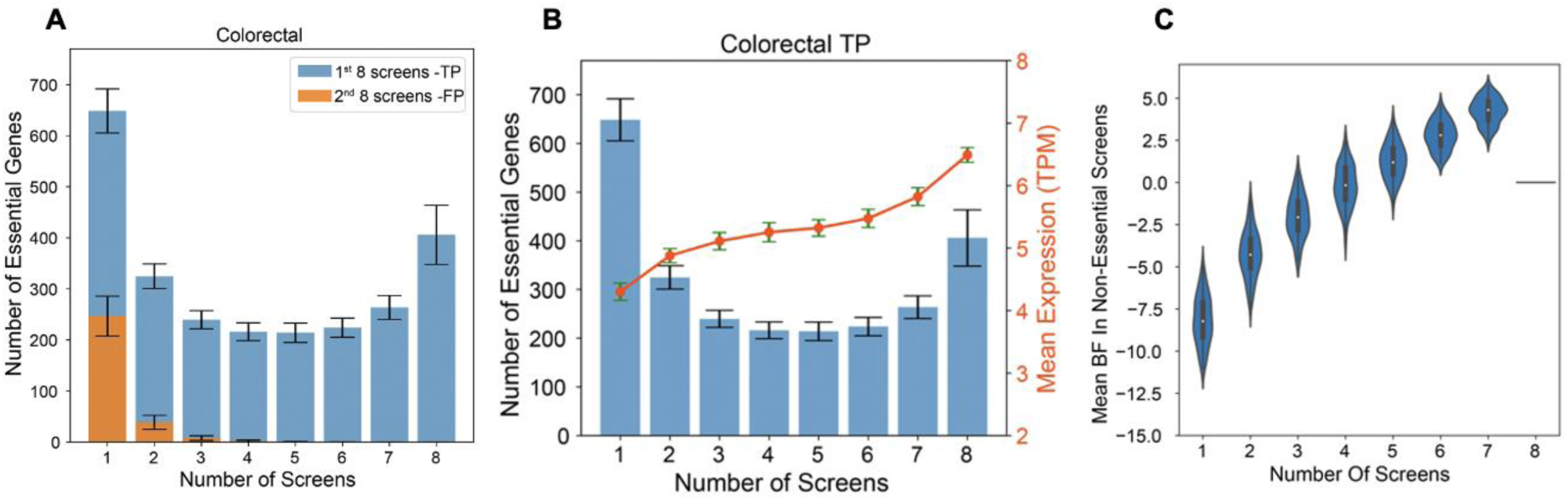
Saturation modeling approach. **A)** Histogram showing the distribution of the number of essential genes and the number of cell lines in which each gene was classified as essential in colorectal cancer cell lines. Blue bars represent the distribution of true positives (TPs) and the orange bars represent the frequency of false positives (FPs). The error bars represent the standard deviation of observed essentials in 100 iterations. **B)** For the TPs in colorectal cancer cell lines depicted in A, the mean mRNA expression (log2(TPM)) of the genes in each bin shows higher expression where more frequently observed essential genes show higher levels of expression. **C)** Violin plot showing the distribution of the essentiality scores (Bayes Factor, BFs) of the TP genes in A and B, in the screens where they were not observed as essential.

Notably, the putative false positives follow the expected distribution: most are detected in only a single screen (**Figure 2A**). We used the distribution of the true and false positives to estimate both binwise and cumulative false discovery rates **(Supplementary Table 2)** by comparing the ratio of putative false positives (subsequent screens) to the total number of hits in each bin (initial screens). For example, if a gene is a hit in only one of eight colorectal screens, we estimate its FDR to be ∼38% (247 first bin FP / 648 first bin hits). We observed that the bin-wise FDR falls to less than 3% for genes that are observed in 3 or more out of 8 screens. Therefore, we determined that genes observed in 3 or more of 8 randomly selected screens represent the high-confidence set of essential genes in a given tissue type and this set includes both genes that are frequently false negatives as well as those that are subtype-specific within a lineage. A complete table of gene frequency of observation by tissue type is presented in **Supplementary Table 3**.

### False negatives and subtype-specific genes

A closer examination of the U-shaped histogram in **Figure 2A** reveals significant implications. Genes that are observed as essential in the intermediate number of screens (∼3 to 6 out of 8) are either false positives that are repeatedly observed, false negatives that are repeatedly missed, or cancer subtype-specific genes that are truly only essential in some cell lines, violating our modeling assumption that the cell lines are identical (in reality, some combination of these three possibilities is more likely). We demonstrate from the hit frequency of the subsequent screens (in orange) that these genes are very unlikely to be false positives. We therefore sought to determine whether these genes are false negatives or background-specific essentials.

First, we examined the mean mRNA expression levels of genes in each bin in the corresponding cell lines that they were detected as essential. We observed a clear trend in which more frequently observed essential genes show higher gene expression **(Figure 2B, right Y-axis)**. In contrast, hits in only one screen show markedly lower expression. In addition, putative false positives from subsequent screens show a similarly lower average gene expression than more frequently observed hits **(Supplementary Figure 5)**.

Next, we examine the essentiality profiles of genes in the screens where the gene is not a hit. This measures whether a gene that is essential (BF>10) in, for example, 5 screens is truly non-essential in the remaining 3 screens with very negative essentiality scores (BF <-10) or whether it falls in the intermediate range near BF=0. **Figure 2C** shows that the more frequently a gene is classified as essential in the initial screens, the higher its average BF in screens where it is not essential. This observation is strongly consistent with false negatives rather than context-specific essential genes.

### High confidence context-essential genes and a newly defined set of core essential genes

After judging that genes observed in 3 or more (out of 8) screens represent the high-confidence set of essential genes in a given tissue type, we identified 992 genes that are essential at that frequency in all 18 lineages we evaluated **(Figure 3A**). Applying these high-confidence essentials to our Daisy Model (T. Hart et al., 2014), we find that each tissue type carries an additional 300- 600 context specific essential genes **(Figure 3A inset),** which make up the petals of the daisy. Interestingly, these additional context-essentials are also widely, but not universally, shared across backgrounds: each lineage has only 3 (central nervous system) to 41 (hematopoietic and lymphoid) genes which are uniquely essential in that lineage. We found many known gene-tissue relationships in this set of unique context essentials. For example, the *SOX10* transcription factor was found to be essential in only skin cells where it plays a major role in the production and function of melanocytes (M. L. Harris et al., 2010; Nonaka et al., 2008). *CTNNB1* and *TCF7L2* are essential only in colorectal cancer cell lines, where activation of the Wnt pathway results in accumulation of *B*-catenin that interacts with and acts as a coactivator for *TCF7L2* that in turn activates downstream genes responsible for colorectal cancer cell survival as well as resistance to chemo-radiotherapy (Albuquerque & Pebre Pereira, 2018; Emons et al., 2017; Murphy et al., 2016). ER^+^ breast cancer cell lines depend specifically on transcription factors *FOXA1* and *GATA3*, which are overexpressed in ER^+^ breast carcinomas (Albergaria et al., 2009; Davis et al., 2016). *E2F1*, which was preferentially essential in pancreatic cancer cells, has been previously shown to regulate both pancreatic B-cell development and cancer growth by increasing the expression of *PDK1* and *PDK3* which results in increased aerobic glycolysis and growth in pancreatic cancers (Denechaud et al., 2017; S. Y. Kim & Rane, 2011; L.-Y. Wang et al., 2016). Overall, our analysis indicates that genes uniquely essential in a particular context are very rare, while hundreds are shared across some but not all tissue types. A complete table of common and context essential genes are listed in **Supplementary Table 4**.

**Figure 3.**
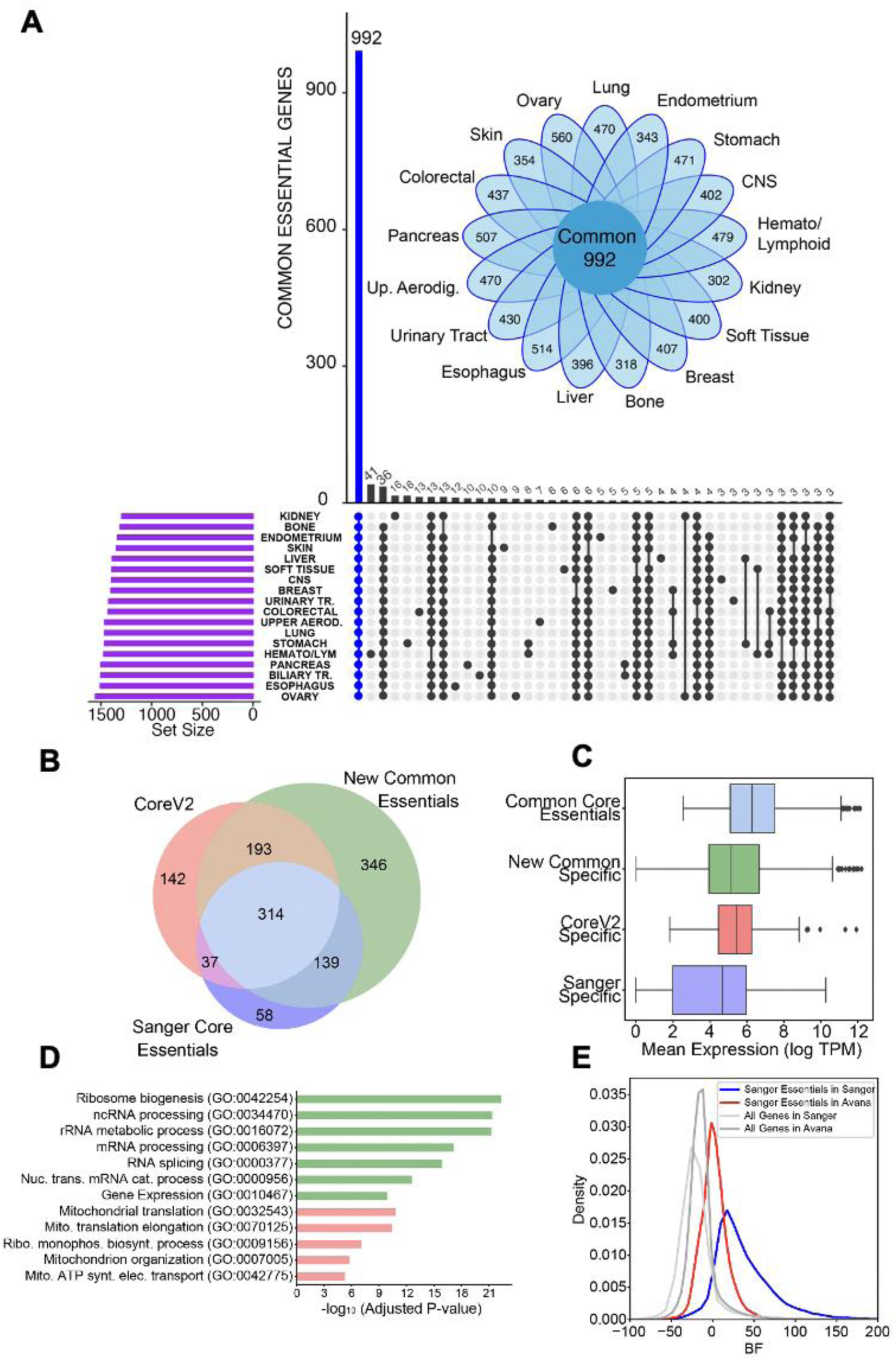
High confidence set of core and context essential genes. **A)** Upset plot showing the intersection of the number of genes between high-confidence essential genes in each tissue type. Inset: Daisy plot showing the relationship between high-confidence context-essential genes and common essential genes. Genes essential in a given tissue type is represented by the petals of the daisy indicating their numbers for each tissue type. The petals overlap to varying degrees, but all tissues share the common set of essential genes (N=954). **B)** Venn diagram comparing common essential genes to previously defined gold standard set of core essential genes. **C)** Box plots showing the mean mRNA (TPM) expression of common core essentials (n=300), genes unique to new common essentials identified here (n=314), Core V2 specific essentials (n=142) and core essential genes specific to the Sanger dataset (n=58). **D)** Gene Ontology (GO) biological process enrichment for core essential genes unique to a specific approach. **E)** Comparison of the distribution of essentiality scores of Sanger specific core essentials in common cell lines between the Avana and Sanger data.

### Comparing common essentials to previous gold standard core essential genes

The common essentials defined in this study include genes that are identified as hits in every tissue type at a frequency of at least 3/8 screens. Therefore, we reasoned that they should define a superset of previously defined gold standard sets of core essential genes. We compared our set of 992 common essential genes to the Core Essential Genes v2 (CoreV2) that we previously defined as a gold standard training set for our BAGEL algorithm (T. Hart et al., 2017), as well as the core essentials defined by the Sanger dataset collected at the Wellcome Trust Sanger Centre (Behan et al., 2019). The newly identified common essentials comprise 507 of 686 (74%) of CoreV2 genes and 453 of 548 (83%) of Sanger core essentials, while 314 genes were common in all approaches (**Figure 3B**). We also examined the gene expression profiles of these core essential genes and observed that the 314 genes at the intersection of the three approaches have higher median gene expression compared to genes that are uniquely core essential in each approach (median log2(TPM) 6.3 vs 5.1 for new common specific genes, 5.4 for CEGV2 and 4.7 for Sanger specific genes; **Figure 3C)**. These observations are consistent with an increased false negative rate among essential genes with moderate levels of mRNA expression. Core essential genes specific to each approach are listed in **Supplementary Table 5.**

Interestingly, the genes unique to each dataset showed a bias reflecting the experimental approach used. Genes unique to common essentials defined in this study are highly enriched for essential processes including ribosome biogenesis and mRNA processing genes (**Figure 3D**). Moreover, genes unique to CoreV2 show a strong bias towards genes encoding subunits of the mitochondrial translation and electronic transport chain. Dependency on oxphos genes is correlated with cell growth rate (Rahman et al., 2020) representing a large source of bias in screen hits, and these genes are likely excluded from the Sanger core essentials for the same reason. We did not observe any strong functional enrichment among Sanger-specific core essentials, compared to the other classes. Among 58 Sanger-specific core essentials, 11 were not targeted in the Avana library and the remainder show intermediate BF scores in the 148 high-performing Avana screens of the same cell lines (**Figure 3E**, red curve). Collectively, these observations are consistent with there being a set of CRISPR library-specific false negatives, as previously reported (Ong et al., 2017), which may be independent of the expression-associated false negatives discussed in this work.

### False negatives are evident in essential protein complexes

In the previous sections, we demonstrated a nontrivial false negative rate in an average genome-wide CRISPR-Cas9 screen (**Figure 1F**). Moreover, we showed that these false negative genes tended to be observed in an intermediate number of screens in our saturation modeling approach. Here, we will highlight the false negatives in essential protein complexes.

Previous studies in model organisms have shown that the individual subunits of protein complexes tend to be either all essential or non-essential (G. T. Hart et al., 2007) excluding cases where there might be functional buffering due to redundancy within the complex members (Ryan et al., 2013). This notion leads to a model where essential protein complexes are comprised of essential genes whereas non-essential complexes contain only non-essential genes. Therefore, in human protein complexes, if a complex has essential functions, we expect to see all obligate members of that complex to be collectively and entirely essential, unless there is functional buffering or alternative subunit usage. To see if the same trend exists in the Avana data, we examined previously defined essential protein complexes to see how consistently the BAGEL algorithm detects individual essential genes. The proteosome is a large, essential protein complex responsible for degrading intracellular proteins (Tanaka, 2009) and we expect this complex to be essential in all cell lines since it carries a core-essential function. In **Figure 4A**, we see the binary essentiality calls (BF>=10) of the members of the 26S proteosome complex as defined in the Comprehensive Resource of Mammalian Protein complexes (CORUM) database (Giurgiu et al., 2019) among colorectal cancer cell lines in the Avana data. We observe that most individual screens are unable to capture all of the individual members of this complex, even though every subunit is detected in multiple cell lines.

**Figure 4.**
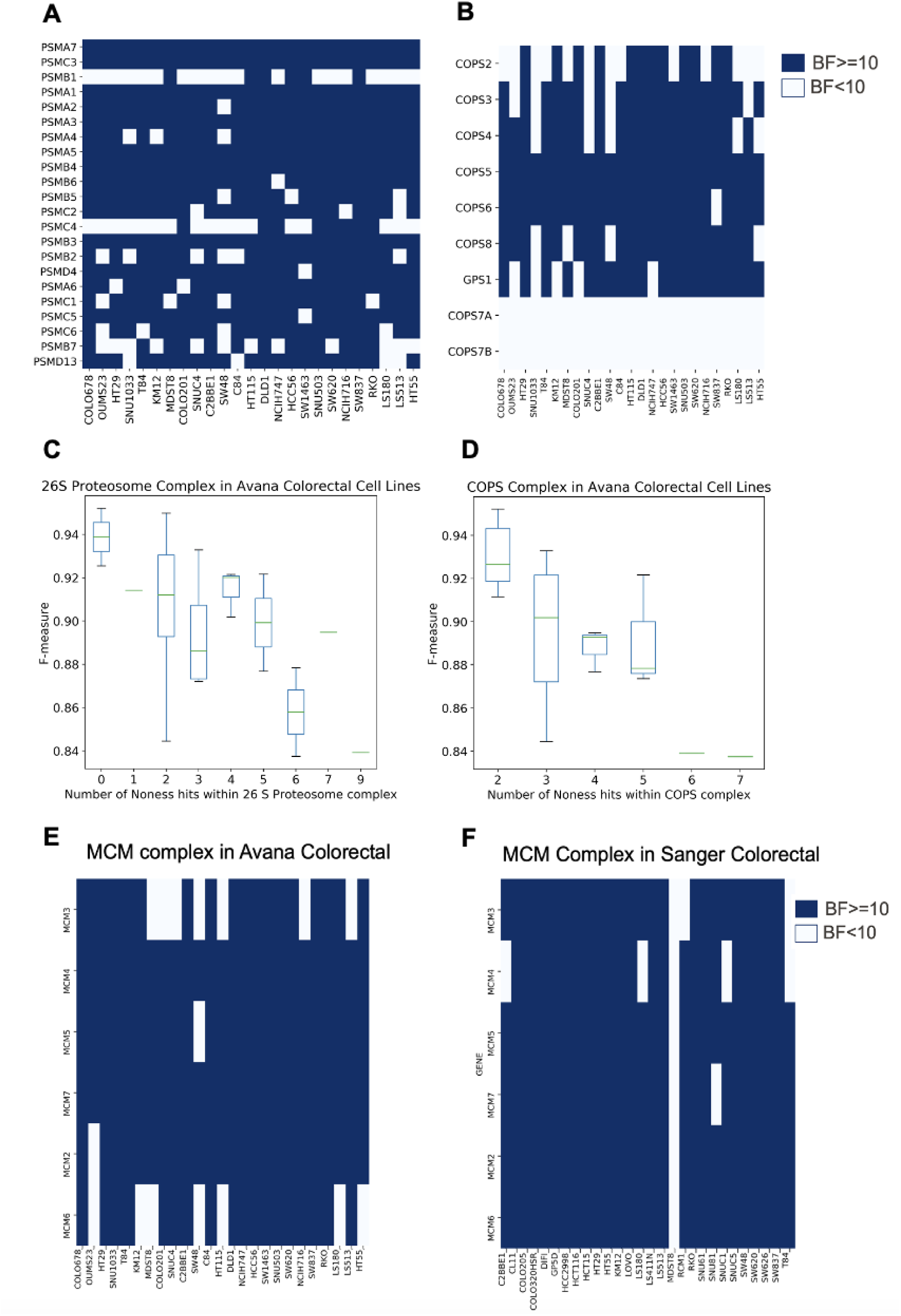
False negative genes in essential protein complexes. Binary essentiality calls of essential protein complexes among colorectal cancer cell lines in the Avana data based on BAGEL derived essentiality scores. Dark blue color indicates essentiality (BF>=10) and white color indicates non-essentiality (BF<10). **A)** Binary essentiality calls of members of 26S proteosome complex. **B)** Binary essentiality calls of members of COP9 signalosome complex among colorectal cancer cell lines in the Avana data based on BAGEL derived essentiality scores. The relationship between screen quality (F-measure) and the number of non- essential hits within essential protein complexes. Boxplots showing the distribution of F- measures of screens versus the number of times a non-essential call was made among members of proteosome complex **(C)** and COPS9 signalosome complex **(D)**. MCM complex is an example of a CRISPR-library specific false negative. While some members of the MCM complex cannot be captured as essential in multiple colorectal screens in the Avana dataset **(E)**, the complex shows more uniform essentiality in Sanger colorectal cell lines **(F)**.

Similarly, in **Figure 4B**, we see the binary essentiality calls in Avana colorectal cancer cell of the members of the essential, evolutionarily conserved COP9 signalosome complex which plays a role in controlling protein ubiquitinylation (Gutierrez et al., 2020). Again, we observe that not all screens can identify members of this complex as essential. In this complex, we don’t observe COPS7A and COPS7B as essential in any screen since, as we and others have previously shown, COPS7A and COPS7B are synthetic lethal paralog pairs and they encode alternate, replaceable subunits of the complex, while the other subunits are irreplaceable and are uniformly essential (De Kegel et al., 2021; Dede et al., 2020).

We first asked whether these false negatives were associated with underlying screen quality. We plotted the F-measure of each screen versus the number of non-essential hits within the complex for both the proteosome and COP9 signalosome complexes. Even though we only considered high-performing screens in our analysis, we still observed a trend where the screens with lower F-measures tended to miss more of these essential genes (**Figure 4C,D**). We further investigated whether these false negatives were library-specific. Comparing Sanger and Avana datasets, we observed that this phenomenon indeed exists as shown for the minichromosome maintenance (MCM) complex, an essential helicase regulating DNA replication in eukaryotic cells (Lei, 2005). While multiple Avana colorectal screens fail to capture members of this essential complex (BF>=10), matched Sanger colorectal screens show more consistent essentiality calls for members of the same complex, indicating that library-specific false negatives also exist (**Figure 4E,F).**

### Sharing BAGEL information across screens: Joint Log Odds of Essentiality

Having shown that CRISPR-Cas9 screens miss expected hits, we sought to develop a method to recapture these false negatives. Our approach extends fundamental concepts from our BAGEL algorithm (see **Methods** for a detailed description). The final output of BAGEL is a log Bayes Factor (BF) which represents the relative statistical confidence of that a gene is a member of one model (essential genes) or another (nonessentials). Based on empirical information from genome-wide CRISPR-Cas9 knockout screens regarding the number of essential genes in a given genome, we estimate the background expectation of gene essentiality in humans as 10%, which yields a log prior ratio ∼-3. When using our BAGEL algorithm, we traditionally use this flat log_2_ prior ratio of -3 during BF calculations for all screens. In order to rescue false negatives, and to differentiate between false negative genes and subtype-specific genes, we employed an alternative method based on the joint analysis of BAGEL results, which we call JLOE (Joint Log Odds of Essentiality). In this approach, instead of using a flat prior ratio for all screens, we update this prior ratio as we gain information from the screens. Gene essentiality is reported as a posterior log odds instead of a Bayes Factor (BF), and binary hit calls are assigned based on a posterior log odds threshold.

Briefly, we evaluated sets of 8 screens at a time in each tissue type. For each gene, we ranked the screens from the highest BF to lowest in each set. Then we calculated the posterior probability of essentiality for that gene in each screen sequentially. For the top ranked screen, we used the initial log prior ratio of -3 and assigned a binary call of essentiality for posterior log(odds) > 7 (corresponding to a posterior Pr(ess) ∼ 99%). For the second-ranked screen, the prior is now based on whether the gene is essential in the top-ranked screen. If so, we use an updated prior ratio derived from the bin-wise FDR calculated from the saturation modeling approach **(Supplementary Table 2)** and again assigned gene essentiality for a joint posterior Log Odds > 7. The process is repeated for the next ranked screen, until all 8 screens are evaluated. Since the bin-wise FDR values are negligible for frequent hits (FDR 0.3% for genes essential in 5 of 8 screens; log prior ratio 8.4), we capped the log prior ratio at 9 to prevent instability. In addition, to ensure that all cell lines are properly sampled, we repeated the sampling process 100 times, selecting 8 screens without replacement in each iteration from every tissue type to minimize bias. The normalized binary calls and posterior log odds from JLOE approach can be found in **Supplementary Tables 6 and 7**, respectively.

### JLOE selectively rescues false negatives

After applying JLOE to all tissue types in the Avana and Sanger datasets, we compared the observations in both datasets to our saturation modeling approach to evaluate differences in the frequency of hits. JLOE resulted in higher frequency of gene essentiality calls, with fewer hits in fewer screens and more hits in more screens – including a large increase in the number of universally essential genes compared to the saturation modeling approach (**Figure 5A** for Avana data). Strikingly, the previously observed trend whereby genes that are hits in a higher number of screens have a higher average BF in the cell lines where they are not hits is completely abrogated by the JLOE approach **(Figure 5B)**, suggesting that JLOE rescues false negatives.

**Figure 5.**
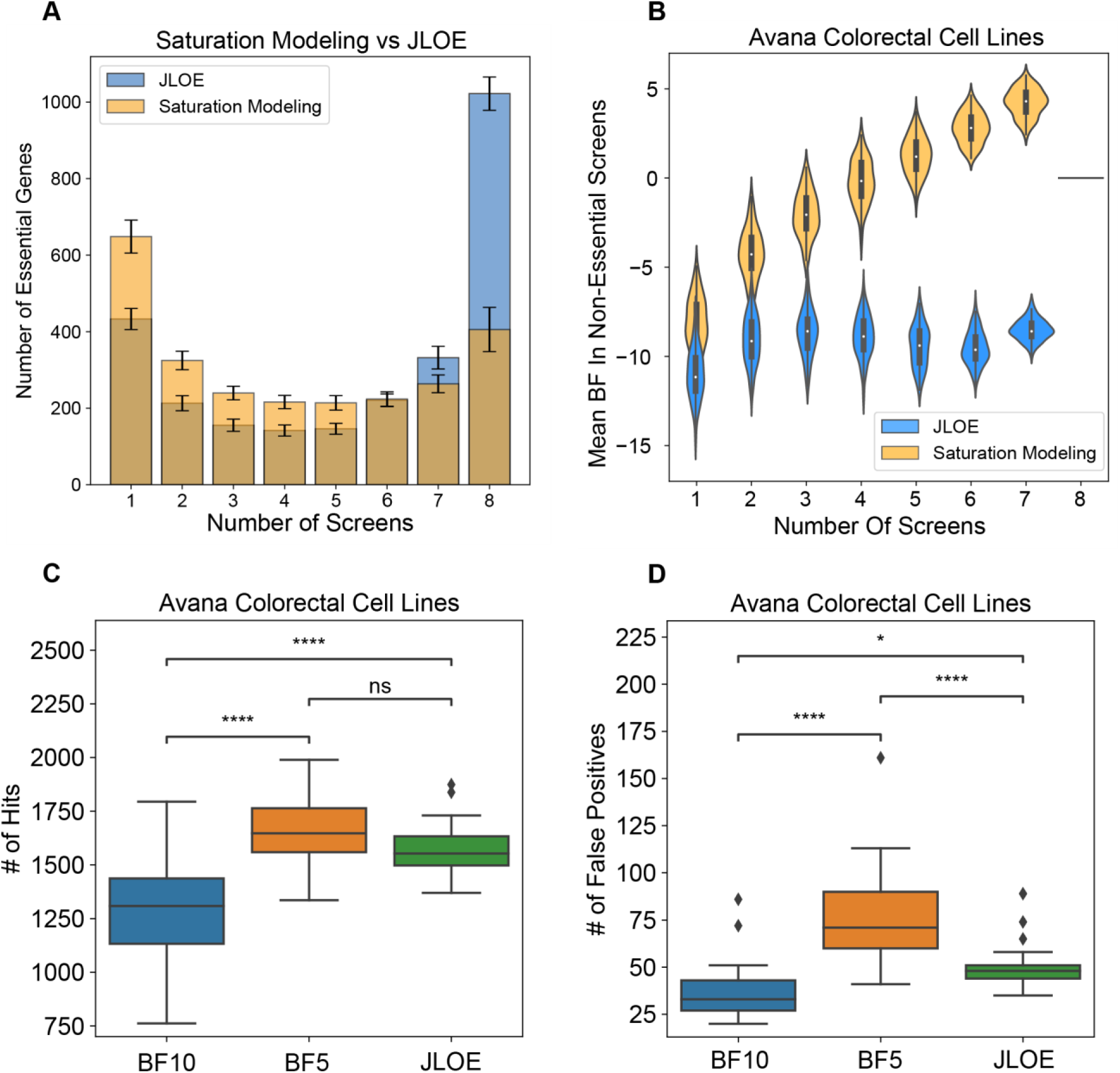
JLOE approach identifies more essential genes in Avana genome-wide CRISPR-Cas9 knockout screens compared to saturation modeling approach. **A)** Comparison of the frequency of essential gene hits in saturation modeling versus JLOE. **B)** Comparison of violin plots showing the distribution of the essentiality scores (Bayes Factor, BFs) of the hits in A, in the screens where they were not observed as essential. JLOE can rescue the false negatives with a less significant increase in false positives compared to assigning binary calls using a lower BF threshold. **C)** The number of essential gene hits using different thresholds (binary calls with BF 10, binary calls with BF 5 and mean binary call of 1 with JLOE across 100 iterations. **D)** The number of false positive genes detected across colorectal cancer cell lines in the Avana data using same thresholds in (C). P-value annotation from t-test ind. samples with Bonferroni correction legend as follows; ns: 0.05 <p, *: 0.01< p <=0.05, **:0.001< p <=0.01, ***: 0.0001 < p <= 0.001.

A concern with boosting scores of marginal results is that false positives are also boosted. If this were the case, JLOE results would be equivalent to simply lowering the BF threshold for binary hit calls. To test this, we compared binary calls at BF>10 (posterior log odds > 7 with uninformative prior) with binary calls at BF>5 (posterior log odds > 2, or Pr(ess) > 80%) and binary calls with JLOE>7 **(Supplementary Table 6)**. Both relaxing the BF threshold and applying the JLOE framework increase the number of hits in Avana colorectal screens compared to the BF 10 baseline (**Figure 5C**). However, JLOE yields a much smaller increase in the number of false positives (defined here as expression log(TPM) < 1) compared to BF>5 (**Figure 5D**), indicating JLOE successfully discriminates marginal essential genes from nonessentials. Strikingly, the mean number of JLOE essentials identified among colorectal cancer cell lines (N=1565) is approximately what we had predicted using our synthetic genome modeling (N=1600), indicating the consistency of our findings using different approaches.

### JLOE increases enrichment of essential processes among high frequency hits

To further confirm that JLOE rescues true essentials, we compared always-essentials (hits in 8/8 screens at BF>10) with JLOE-classified false negatives (hits in < 8/8 screens at BF>10 but 8/8 screens after JLOE) in both Avana and Sanger screens in colorectal cell lines (n=25 each). Always-essentials in Avana (n=359 genes) and Sanger (n=403 genes) screens show significant overlap (**Figure 6A** – Jaccard coefficient 0.339), but more disjoint sets of false negatives (**Figure 6B**; Jaccard coefficient 0.182). Combining the always-essentials with the JLOE-rescued false negatives yields not only more genes (Avana, n=937; Sanger, n=839), but also better concordance between the two sets (intersection n=610 genes, Jaccard coefficient =0.523; **Figure 6C**). Moreover, boosting always-essentials with JLOE-rescued false negatives improves gene annotation enrichment statistics for core essential processes including transcription, translation, and cell cycle regulation in both Avana (**Figure 6D**) and Sanger (**Figure 6E**) screens. Overall, these results show that the JLOE approach can rescue false negative genes from essential pathways and improves the concordance between the Avana and Sanger datasets.

**Figure 6.**
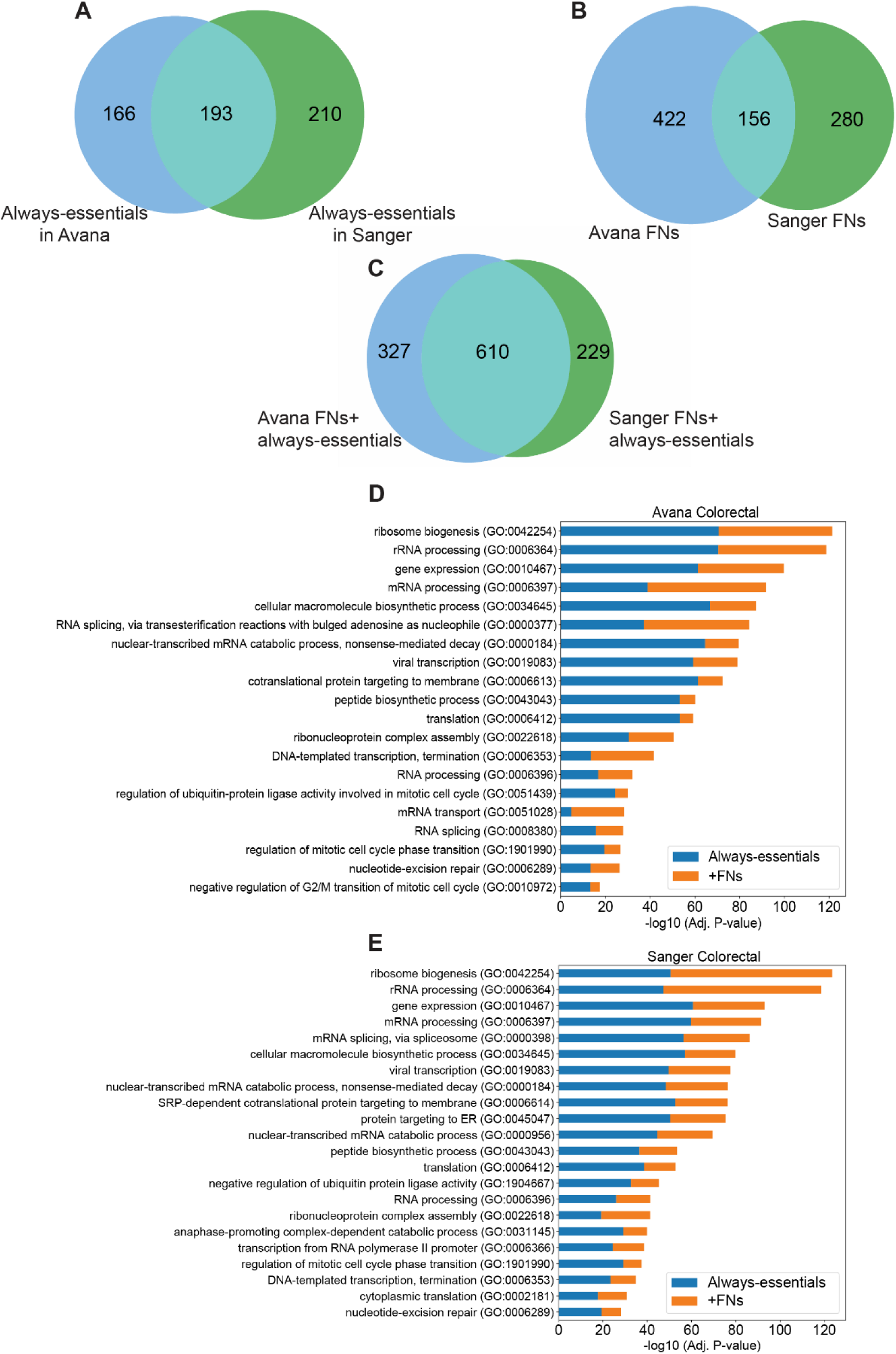
JLOE increases the concordance between Avana and Sanger datasets. **A)** The overlap of always-essential genes (with frequency observations of 8/8 in saturation modeling approach among colorectal cancer cell lines in Avana and Sanger datasets. **B)** Venn- diagram showing the overlap of the false negative genes among colorectal cancer cell lines in Avana and Sanger datasets. **C)** Combination of false negative genes with always-essential hits in saturation modeling approach in both datasets. The recovered false negatives increase the enrichment of essential biological processes in colorectal cell lines among high frequency hits defined in saturation modeling approach in **(D)** Avana colorectal cell lines and **(E)** Sanger colorectal cell lines. Blue bars indicate the enrichment among always-essentials and orange bars depict the increase in enrichment with the addition of false negative genes with JLOE.

## Discussion

The vast amount of publicly available human cancer cell line data generated by the international Cancer Dependency Map initiative offers an opportunity to re-evaluate the assumptions under which these genome-scale experiments have been carried out. One such assumption is that CRISPR screens are saturating and that a well-designed single whole genome CRISPR-Cas9 knockout screen can detect virtually all of the essential genes in that background. Although library -specific biases have been identified (Ong et al., 2017) and copy number amplifications are known induce false positives, our knowledge of the systematic biases in CRISPR screens is likely still incomplete. Simply put, the true positive, false positive and false negative rates in an average genome-wide CRISPR-Cas9 knockout screen are fuzzy at best.

Here, we describe modeling approaches to estimate the true positive rate, false positive rate, and false negative rate from a typical genome-scale CRISPR screen. Importantly, these analyses established a baseline for the expected number of essential genes and the FDR in an average screen, with the broad definition of “essential gene” being one whose knockout fitness defect causes the mutant to be outcompeted by cells growing at wildtype proliferation rates. Our modeling suggests that a typical cell expresses 1,600-1,900 essential genes, but a single knockout screen only detects ∼80% of these, and multiple screens are required to saturate the essential genes of any tissue or tumor subtype. We show that hits among highly expressed genes are often replicated but false negatives are more prevalent among genes expressed at moderate levels. These characteristics may limit our ability to identify meaningful differentially essential genes, in particular using isogenic cell lines to identify synthetic lethal interactions, since it suggests numerous replicates need to be screened in order to confidently discriminate cell-line- specific hits from false negatives and false positives. This knowledge is critical for evaluating the significance of findings from current experiments as well as for the design of future experiments for gene essentiality and synthetic lethality.

Another implication of the false negative rate is that most tissue types carry a larger number of overlapping essential genes than had been previously recognized. We identify nearly a thousand genes that are observed across all lineages that are frequent flyers in the Avana screens. This improved new set of core essential genes are enriched in core essential processes of the cells and, like the previous set, can be used as a metric for the quality control of genome-wide CRISPR- Cas9 knockout screens. In contrast, we find only 300-600 tissue-specific genes per lineage, with the overwhelming majority of genes showing overlap between related tissues. In fact, each tissue only carries at most a few dozen tissue-unique genes, and these are highly enriched for lineage- specific transcription factors.

Genes we classify as likely false negatives can be recovered though an approach we call the Joint Log Odds of Essentiality, or JLOE, which uses the Bayesian framework of BAGEL to improve analysis across multiple cell line screens. An unexpected, and unexplained, finding of our study is that these false negatives exhibit more moderate mRNA expression levels than previously described always-essential genes. This opens new questions about the biological relationship between expression level and the severity of loss of function phenotype, on one hand, and concerns about the effects of chromatin state on the relative efficacy of the CRISPR/Cas9 knockout paradigm on the other.

## Methods

### Essentiality Data Generation

The raw read count file of genome-wide CRISPR pooled library screens for 769 cell lines using Avana library (Broad DepMap, 2020b; Meyers et al., 2017) (Broad Institute’s DepMap Project 20Q2 release) was downloaded from the data depository at https://depmap.org/portal/. The dataset was filtered in order to keep only the protein-coding genes for downstream analysis and the gene names were updated using Human Genome Organization (HUGO) Gene Nomenclature Committee (HGNC) (Braschi et al., 2019) and Consensus Coding Sequence (CCDS) (Farrell et al., 2014) databases. sgRNAs targeting multiple genes were discarded to avoid genetic interaction effects. The updated raw read counts were processed with the CRISPRcleanR (Iorio et al., 2018) algorithm to correct for gene- independent copy number induced fitness effects and calculate fold change. CRISPRcleanR uses a circular binary segmentation algorithm which was previously used during the analysis for array-based comparative genomic hybridization assay (Olshen et al., 2004; Venkatraman & Olshen, 2007) and applies it to genome-wide CRISPR screens. For each individual chromosome, CRISPRcleanR detects regions targeted by multiple sgRNAs with reasonably equal log fold changes (logFCs). If the sgRNAs in these regions target a low number of unique genes, this would indicate that the phenotype arises due to gene-independent effects and therefore the logFCs are corrected through mean or median based centering depending on the presence of outliers (Iorio et al., 2018). CRISPRcleanR processed fold changes of each cell line were analyzed through updated BAGEL2 algorithm (E. Kim & Hart, 2021) (https://github.com/hart-lab/bagel). Compared to the previous published version of BAGEL (T. Hart & Moffat, 2016), the updated version of BAGEL employs a linear regression model to interpolate outliers and uses 10-fold cross validation for data sampling. BAGEL is a Bayesian classifier that is trained using previously defined gold standard reference sets of core-essential and nonessential gene sets. BAGEL estimates the distribution of fold changes of all gRNAs targeting all genes in either the essential or nonessential training sets and then it calculates the log likelihood of uncharacterized sgRNAs belonging to either the essential or nonessential distributions and gives a log Bayes Factor (BF) as the final output. Essentiality of each gene was measured as BF, which reflects the relative statistical confidence of gene essentiality based on gold standard reference sets of 681 core essential genes and 927 nonessential genes (T. Hart et al., 2014, 2017). Positive BF indicates essential genes whereas negative BF indicates non-essential genes. The list of gold standard core-essential genes and nonessential genes can be found in the same repository as the BAGEL v2 software. The qualities of each screen was evaluated through the “precision-recall” function of BAGEL and the F-measure, which is the harmonic mean of the precision recall, was calculated for each screen at BF of 5. Finally, 659 cell lines for the Avana dataset were selected for downstream analysis by an F- measure threshold of 0.80 to prevent noise from marginal quality of the screens.

### Cumulative Essentials Analysis

A cumulative analysis approach was used to evaluate and display the cumulative distribution of essential genes and calculate the total number of true essentials (true positives) as well as the error rate (FDR) in an average CRISPR- Cas9 knockout screen. This approach is based on the principle described previously (T. Hart et al., 2014) that if you screen a sufficient number of cell lines with no false positives, you would expect to see a cumulative essential genes observations plot that flattens out with a slope of zero when the total number of essential genes is reached.

However, in reality, if you have some FDR or heterogeneity in the population of cell lines being screened, the slope of the cumulative essential plot would stay positive. This is because the true hits get saturated but in addition, it is likely that false positives as well as cell-line specific context essential genes will also be captured and be added on in each consecutive screen.

To perform the cumulative essentials analysis for the screens in the DepMap project, herein referred to as the Avana data, the BFs for all genes across all cell lines was constructed in a separate matrix and the cumulative essentials plot was modeled in any sets of 20 screens from the dataset. First, an initial set of 20 cell lines were sampled without replacement and the essential genes in the first screen out of 20 were identified with a BF of greater than or equal to 10. Then the subsequent screen was evaluated to obtain the essential genes in that screen and the newly discovered essential genes that were not identified in the prior screen were added to the list of essential gene hits to obtain a cumulative essential gene list. The process was repeated for all 20 cell lines to capture all cumulative essentials in the 20 screens. This random sampling process was repeated 100 times to prevent bias in the modeling process. The resulting mean cumulative essentials curves were plotted with standard deviations reflecting the variability of cumulative essential genes observed in each of 100 iterations. Moreover, for each iteration, the newly discovered essential genes in each set of 20 screens were also identified and displayed on the same plots.

### Synthetic Genome Modeling Approach

To estimate the number of essential genes per screen as well as the average error rate, in silico simulations of synthetic screens were conducted. First, for each dataset, a synthetic genome was constructed with the number of genes assayed in that library (i.e. for Avana library N=18,111). For a given genome with N number of genes in it, there is a number of true essential genes; represented by n, and number of non-essential genes (N-n). Then, the precision (1 - FDR) of the assay is represented by the ratio of true positives to that of the total number of hits. We defined a range of thresholds for false discovery rate to test in our model from 1-15%. We then randomly sampled this synthetic genome with a screen with randomly drawn 1287 hits for the Avana dataset (the mean number of essential gene hits at BF>=10 across all Avana screens) from the essential (n) and non-essential (N-n) populations based on the defined FDR in the simulation (e.g. for 10% FDR, 129 nonessentials and 1,158 essentials were randomly selected). We observed the cumulative essential genes across 8 iterations since we estimated that sampling 8 screens was a good estimate of observing the trend in screen saturation in the Avana data. At the same time, we constructed a mean cumulative essentials curve determined from the Avana data for any 8 screens using bootstrapping for 100 iterations. After running the simulations for a range of different number of essential genes (n) and FDR values, cumulative observation curves were plotted for each simulation. The root mean squared error for every synthetic screen was calculated by evaluating the difference between the observed values and the cumulative essentials curve determined from the Avana data.

Since the Avana data contains cell lines arising from multiple tissue types, it is possible that some of the tissue specific essential genes would be wrongly classified as false positives. Therefore, in order to minimize this effect, the synthetic genome modeling approach was repeated for individual tissue types which were represented by at least 16 high-quality cell lines (as defined by F-measure of greater than or equal to 0.8) using the same parameters described above. All of the analyses were performed using Python version 3.6 using multiple packages including pandas (Reback et al., 2020), NumPy (C. R. Harris et al., 2020), the sklearn.metrics, sklearn.utils, resample modules in SciKits (Buitinck et al., 2013), Matplotlib (Hunter, 2007), seaborn (Waskom et al., 2020), and scipy (SciPy 1.0 Contributors et al., 2020).

### Saturation Modeling Approach and Identification of High-Confidence Essential Genes

While the in-silico simulations with synthetic genome model enabled an estimation of the average number of essential genes per screen with an upper limit for the FDR, they didn’t give information about which genes were truly essential. To distinguish truly essential genes from false positives, we identified the tissue types in the Avana data that were well represented (n>=16 screens). 18 tissue types in the Avana fit our criteria and for these tissue types, the frequency of essential gene observations in any 8 screens was evaluated separately for each dataset. For each tissue type, a set of 8 initial screens were selected, the cumulative essential genes in this set were identified and the frequency of essentiality observations was plotted for this set. For the modeling, it was assumed that if these first 8 screens have reached saturation (to approximate for our model), then the subsequent hits in the next set of screens would give us false positives. Therefore, a subsequent set of 8 screens from the same tissue type were randomly selected without replacement and the newly discovered cumulative essential genes that were not present in the initial screens were identified to model the frequency distribution of the false positives. This process was repeated 100 times for each tissue type and the resulting distribution of essential gene counts in these screens were plotted, which were used to calculate the bin-wise FDR. For the 100 iterations that were performed per tissue type, genes observed as essential in at least 3 screens on an average out of 8 were considered as high confidence essential genes in that tissue. Finally, we assessed how many genes were captured as essential in common in all tissues to find the set of “common” essential genes (n=954) and context essentials in each tissue type. We used the UpsetR package (Conway et al., 2017) in R programming language to visualize the set of intersections of essential genes across all tissue types for both of the datasets.

### Expression Data and Analysis

The log2 transformed RNA-seq TPM expression data was utilized from DepMap Data Portal for the 2020Q2 release (Broad DepMap, 2020) for this analysis. The mean TPM expression levels were calculated for all genes in each bin in their corresponding cell lines in which the gene was a hit. The process was repeated for all the bins from the histogram of the frequency of essentiality observations of the initial set of 8 screens representing the true positives and the subsequent set of 8 screens representing false positives.

### Essentiality Profiles of Hits in Non-Essential Screens

In this analysis, we evaluated the mean essentiality scores of genes representing true positives in the screens that they were not observed as essential in. For each iteration, we determined the screens in which genes in each bin had a BF <10 and calculated a mean essentiality score for each gene in those screens. We then visualized the distribution of mean essentiality score observations of genes in each bin in the non-essential screens as a violin plot using the Seaborn package (Waskom et al., 2020).

### Joint Log Odds of Essentiality (JLOE) Approach

In order to explain the methodology behind JLOE, we need to re-visit some details about our BAGEL algorithm. The output of BAGEL is the Bayes factor (BF), the statistical representation of which can be displayed with the equation below:

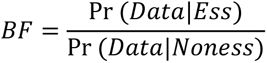

This equation can be converted to a posterior log odds as follows by multiplying by an appropriate ratio of priors:

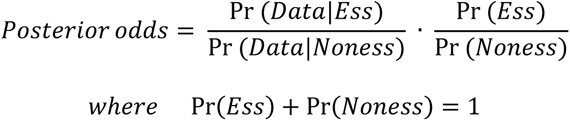

Performing log transformation, we get:

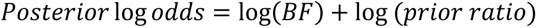

Based on empirical information from genome-wide CRISPR-Cas9 knockout screens regarding the number of essential genes in a given genome, BAGEL considers Pr(*Ess*) = 0.1 (ie. The background expectation of gene essentiality in humans is 10%), which yields Pr(*Noness*) = 0.9 and therefore we have:

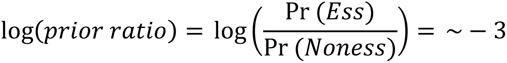

The relationship between the posterior probability of essentiality and BFs can be seen in Supp Fig. 1C, which shows that BF of 10 corresponds to a posterior probability of essentiality of ∼99% and that is why we use this strict threshold for identifying essential genes. During BF calculations with our BAGEL algorithm, we traditionally use this flat prior ratio of -3 for all screens. However, JLOE relies on the notion that this prior ratio gets updated as we gain more observations from the available data. The final outcome is a binary essentiality call and JLOE score based on posterior log odds ratio for each gene instead of a BF.

First, we revisited our saturation modeling approach, where we evaluated sets of 8 screens at a time in each tissue type. Then we calculated the posterior probability of essentiality for that gene in each screen sequentially. For each gene, we ranked the screens from the highest BF to lowest. Next, for the top ranked screen, we used the initial log prior ratio of -3 and assigned a binary call of essentiality for posterior log(odds) >7 (corresponding to a posterior Pr(ess) of 99%). For the second-ranked screen, the prior is now based on whether the gene is essential in the top-ranked screen. If so, we use an updated prior ratio derived from the bin-wise FDR calculated from the saturation modeling approach. For FDR ∼ 50%, the log prior ratio is approximately zero:

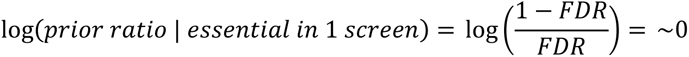

Again, a joint posterior Log Odds > 7 indicates gene essentiality. The process is repeated for the next ranked screen, until all 8 screens are evaluated. Since the bin-wise FDR values are negligible for frequent hits (FDR 0.3% for genes essential in 5 of 8 screens; log prior ratio 8.4), we capped the log prior ratio at 9 to prevent instability. In addition, to ensure that all cell lines are properly sampled, we repeated the sampling process 100 times, selecting 8 screens without replacement in each iteration from every tissue type to minimize bias. For every tissue type, we recorded all binary calls and joint posterior Log Odds for all genes in 100 iterations as well as the cell lines that were sampled in each iteration. Finally, we converted these tissue level scores (frequency of observation out of 8 screens) to cell line level assignments by calculating the total number of times a binary call of essentiality was made for each gene in each cell line across 100 iterations with JLOE and then normalizing by how many times that cell line was chosen in our 100 iterations. With this method, we obtained normalized binary essentiality calls at a cell line level that ranged from 0 to 1 for each gene as well as normalized JLOE scores. We provide these tables across all tissue types in **Supplementary Tables 6 and 7**, respectively.

### Identification of false negative genes in tissues

Mean frequency observations out of 8 screens with JLOE approach were compared to those from the saturation modeling approach. The observations were binned from each method and hits in < 8/8 screens at BF>10 with saturation modeling approach but 8/8 screens after JLOE in both Avana and Sanger screens were classified as false-negative genes across all tissue types.

## Supplementary Figures

**Supplemental Figure 1.**
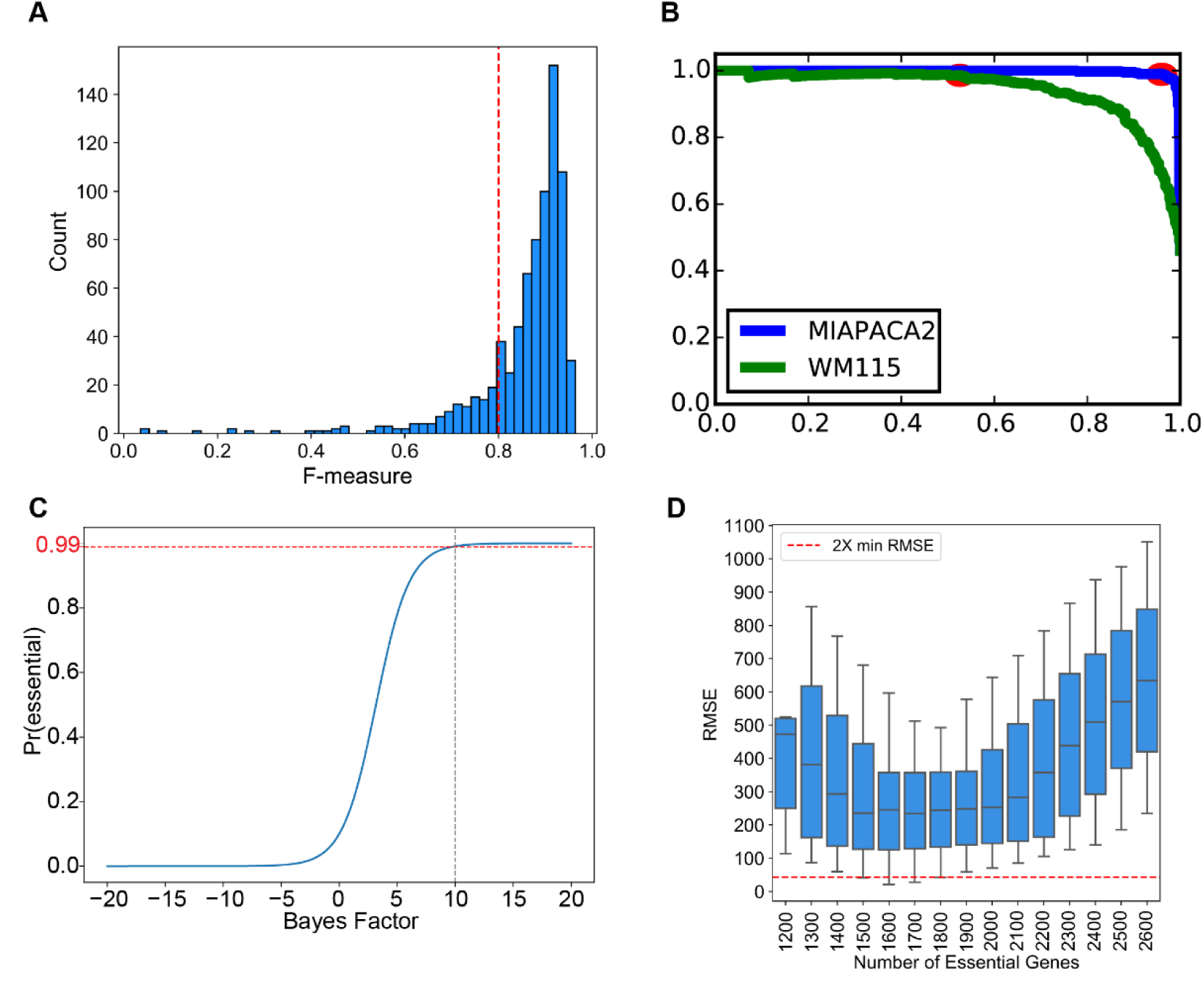
**A)** the distribution of the F-measures of 769 Avana Screens. The red dashed line indicates F-measure of 0.8 and screens with F-measure >0.8 were considered as high-performing and were retained for downstream analyses. **B)** Precision-recall curve specifying an example of a good performing screen (MIAPACA2, in blue) and a bad performing screen (WM115, in green). For all Avana screens, precision-recall curves were calculated using the reference gold standard sets of essential and non-essential genes and the point on the precision-recall curve for each screen where the BF crossed 5 (red points) were identified and the F-measure of each screen was calculated at that point. **C)** Bayes factor of 10 represents a strict threshold corresponding to a posterior probability of gene essentiality of ∼99% **D)** The distribution of the root mean squared deviation (RMSE) values for each simulation reveals a range of models with RMSE < 2xRMSE_min_ indicated by the red dashed line.

**Supplemental Figure 2.**
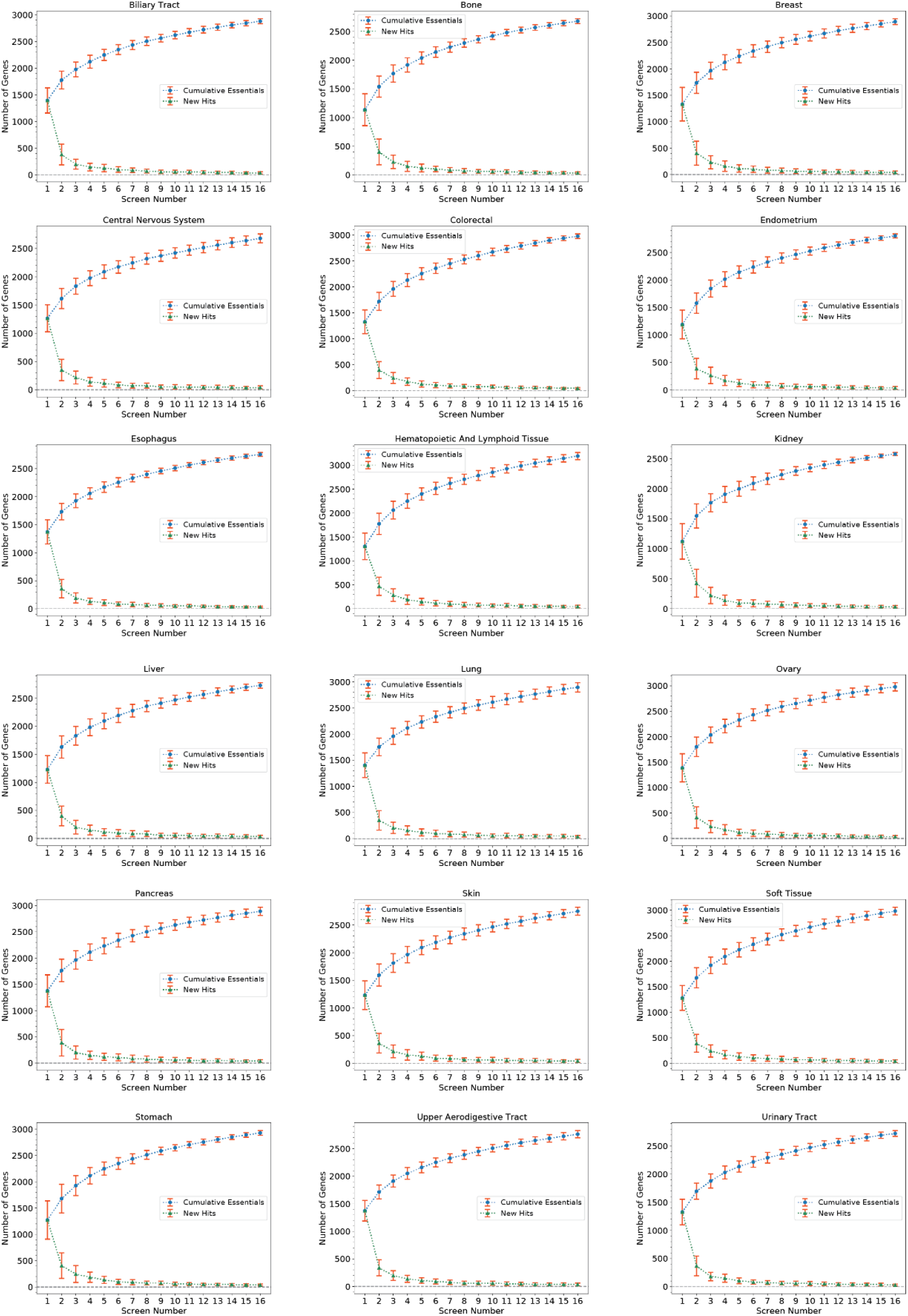
The cumulative essentials curves for lineages represented by more than or equal to 16 high-quality screens. Sets of 16 cell lines were randomly selected without replacement from all screens and the number of cumulative essential genes with BF>=10 in each consecutive screen were plotted in blue with the error bars indicating the standard deviation of cumulative essential gene observations across 100 iterations. The number of newly discovered essential genes in each consecutive screen was also plotted in green

**Supplemental Figure 3.**
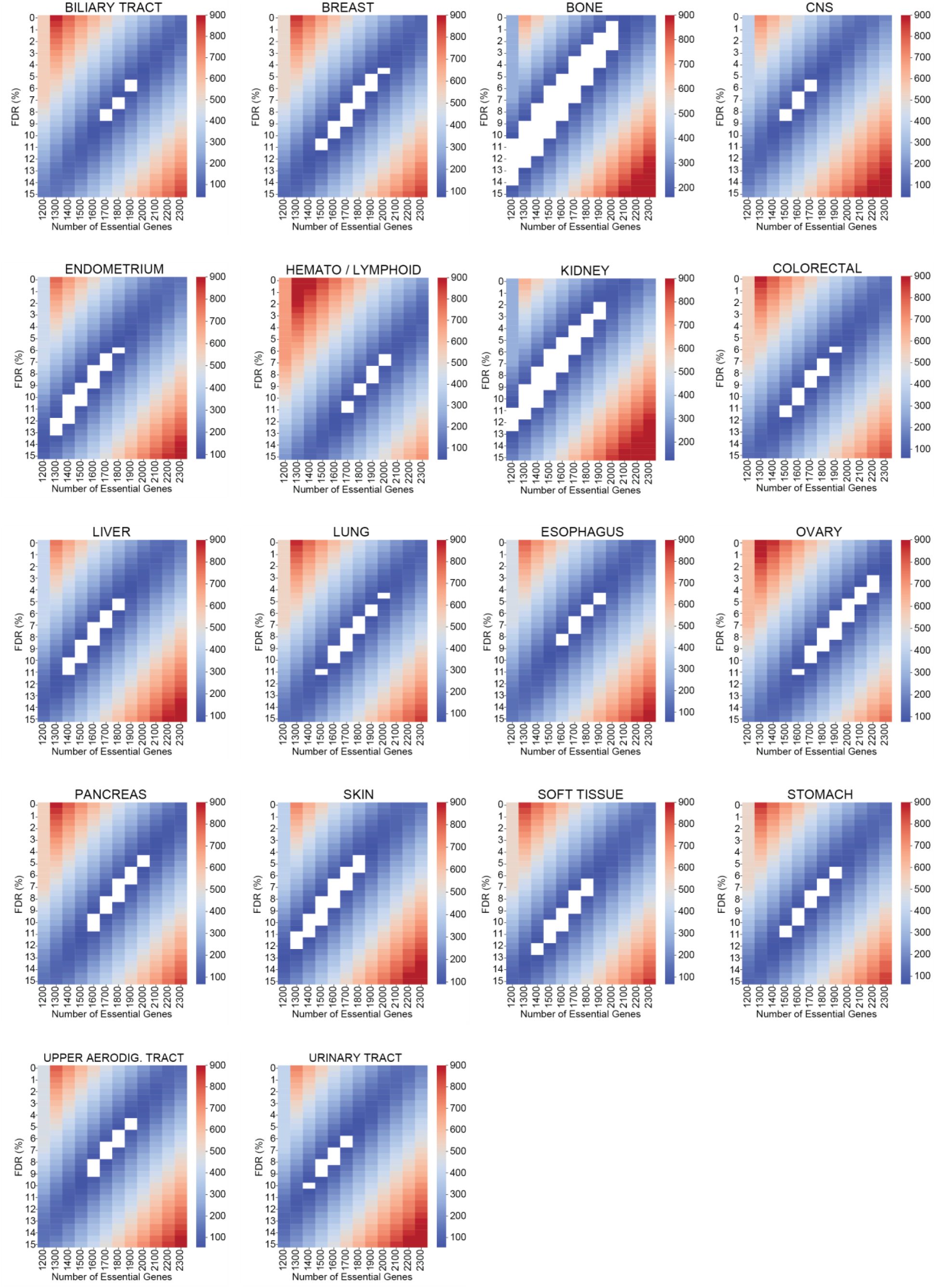
Synthetic genome modeling, applied to each tissue type, estimates the number of essential genes and false discovery rate (FDR) per tissue. Heatmaps showing the root mean squared deviation (RMSE for the models versus the FDR and the number of essential genes in each simulation. The white boxes indicate models with RMSE <2 x RMSE_min_.

**Supplemental Figure 4.**
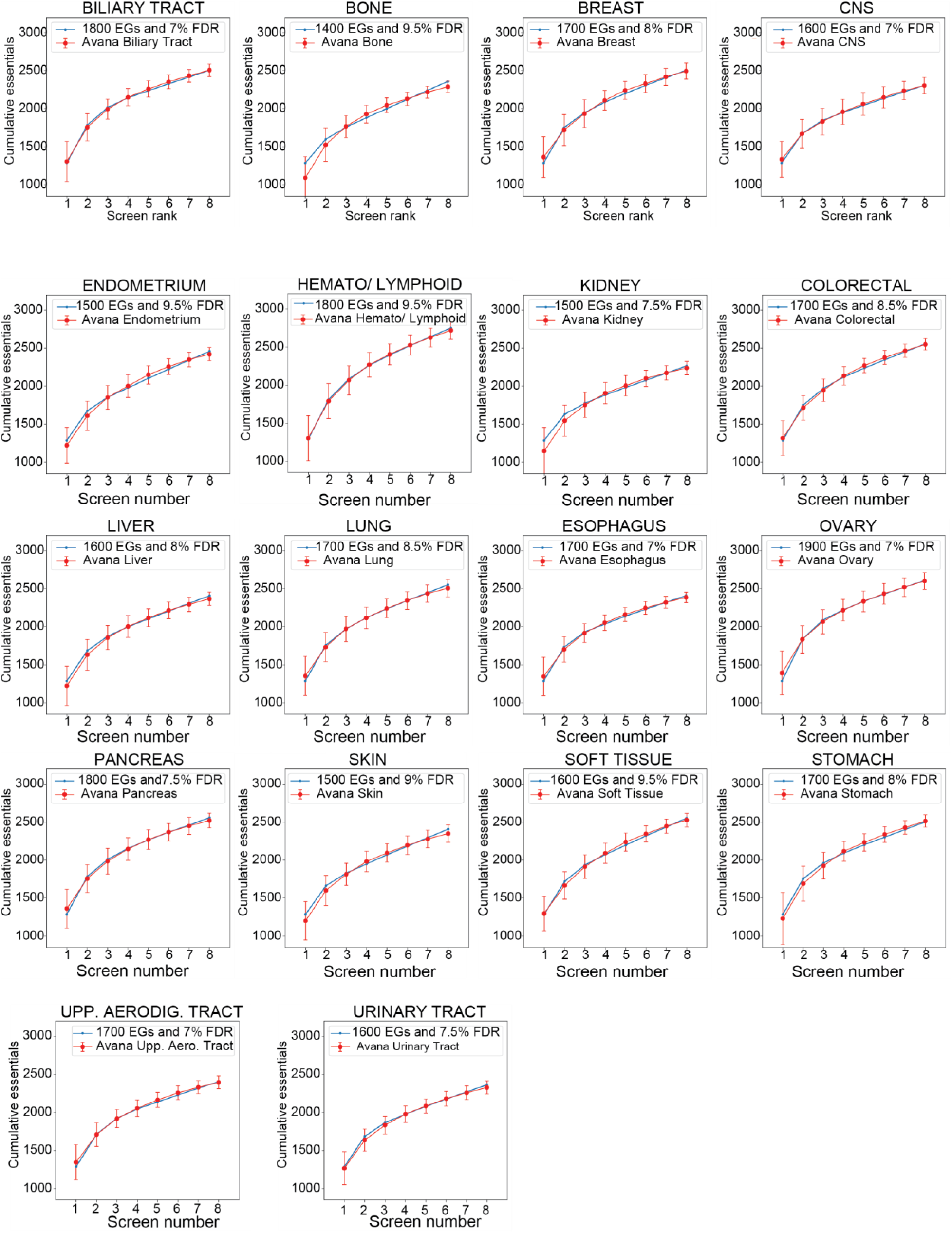
The best fitting models from the synthetic genome modeling approach for individual tissue types. The cumulative essentials curves were plotted for the best fitting model indicated by the blue lines and their fit to the Avana data in their corresponding tissue types (cumulative essential genes across sets of 8 call lines randomly selected without replacement from all available screens in that tissue type for 100 iterations) is shown in red.

**Supplemental Figure 5.**
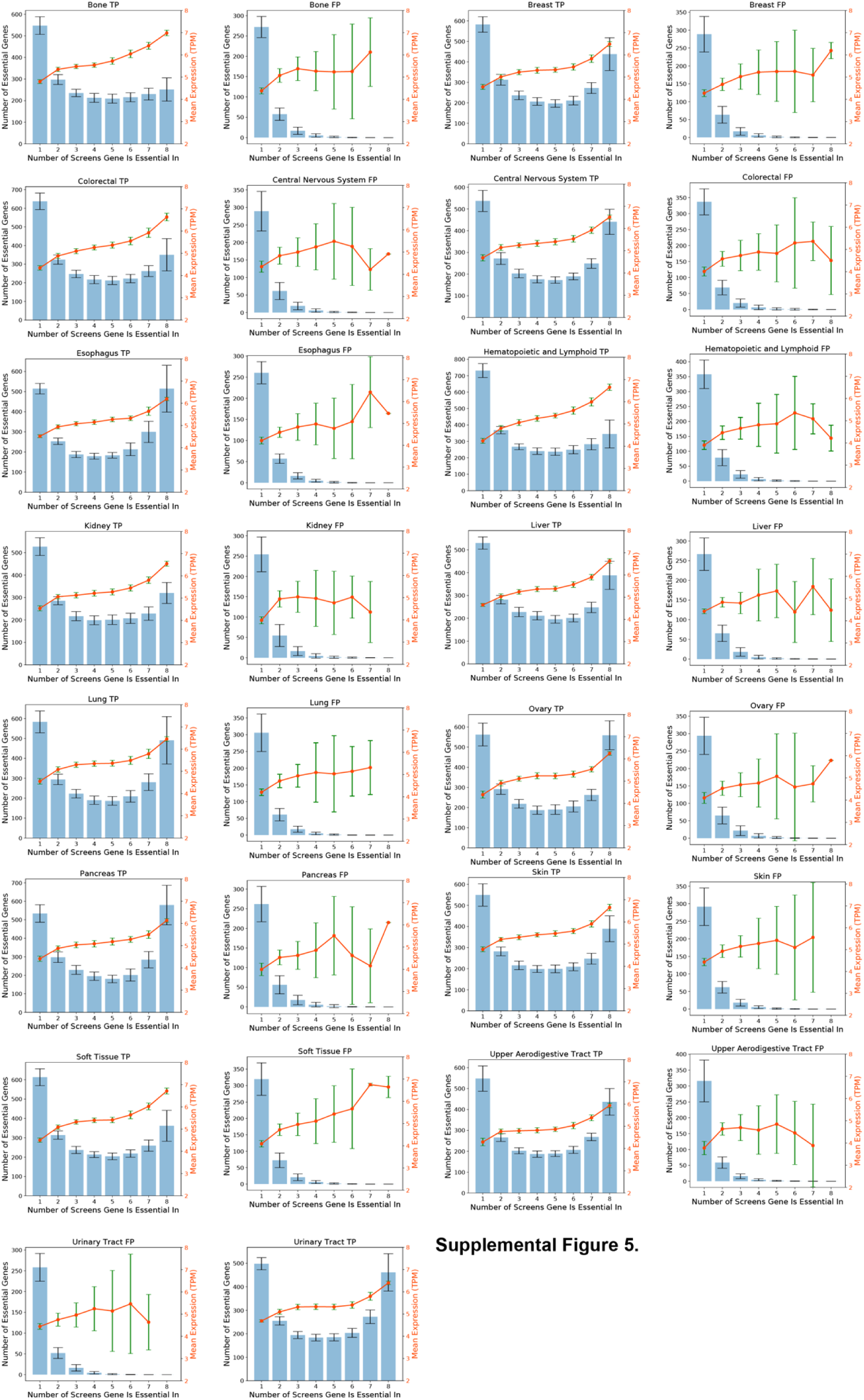
The number of genes in each bin and the mean mRNA expression (TPM) of the genes (indicated by the secondary Y-axis in orange) in corresponding bins for each tissue type for putative true positives (TPs) and false positives (FPs). Error bars indicate the standard deviation of expression of genes in each bin.

## Supplementary Table Legends

Supplementary Table 1. Bayes Factors for the 659 cell lines used in this study with F-measure above 0.80 post CrisprCleanR processing.

Supplementary Table 2. Table of binwise false discovery rates across 100 iterations for the tissue types investigated in this study.

Supplementary Table 3. Gene frequency observations out of 8 screens across 100 iterations by tissue type.

Supplementary Table 4. Table of 992 common essential genes and high confidence context essential genes in each tissue type.

Supplementary Table 5. Common essential and core essential genes unique to each approach among previously defined core essential genes.

Supplementary Table 6. Table of normalized binary essentiality calls from JLOE.

Supplementary Table 7. Table of normalized joint posterior Log Odds from JLOE.

## Acknowledgments

MD and TH were supported by NIGMS grant R35GM130119. MD was supported by a Schissler Foundation fellowship. TH is a CPRIT Scholar in Cancer Research, an Andrew Sabin Family Fellow, and is additionally supported by MD Anderson Cancer Center Support Grant P30 CA016672.

